# Insular cortex sub-region-dependent distribution pattern of α-synuclein immunoreactivity in Parkinson’s disease and dementia with Lewy bodies

**DOI:** 10.1101/156984

**Authors:** Yasmine Y. Fathy, Frank Jan de Jong, Anne-Marie van Dam, Annemieke J.M. Rozemuller, Wilma D.J. van de Berg

## Abstract

The insular cortex is a heterogeneous and widely connected brain region. It plays a role in autonomic, cognitive, emotional and somatosensory functions. Its complex and unique cytoarchitecture includes a periallocortical agranular, pro-isocortical dysgranular, and isocortical granular sub-regions. In Parkinson’s disease (PD), the insula shows α-synuclein inclusions in advanced stages of the disease and its atrophy correlates with cognitive deficits. However, little is known regarding its regional neuropathological characteristics and vulnerability in Lewy body diseases. The aim of this study is to assess the distribution pattern of α-synuclein pathology in the insular sub-regions and the selective vulnerability of its different cell types in PD and dementia with Lewy bodies (DLB). Human post-mortem insular tissues from 10 donors with incidental Lewy body disease (iLBD), PD, DLB, and age-matched controls were immunostained for α-synuclein and glial fibrillary acid protein (GFAP). Results showed that a decreasing gradient of α-synuclein pathology was present from agranular to granular sub-regions in iLBD, PD and PD with dementia (PDD) donors. The agranular insula was heavily inflicted, revealing various α-synuclein immunoreactive morphological structures, predominantly Lewy neurites (LNs), and astroglial synucleinopathy. While dysgranular and granular sub-regions showed a decreasing gradient of inclusions and more Lewy bodies (LBs) in deeper layers. In DLB, this gradient was less pronounced and severe pathology was observed in the granular insula compared to PDD and regardless of disease stage. Protoplasmic astrocytes showed α-synuclein inclusions and severe degenerative changes increasing with disease severity. While few von Economo neurons (VENs) in the fronto-insular region revealed inclusions, particularly in PDD patients. Our study reports novel findings on the differential involvement of the insular sub-regions in PD and particular involvement of the agranular sub-region, VENs and astrocytes. Thus, the differential cellular architecture of the insular sub-regions portrays the topographic variation and vulnerability to α-synuclein pathology in Lewy body diseases.

## Introduction

Parkinson’s disease (PD) is the second most common neurodegenerative disease, mainly characterized by motor symptoms due to the death of dopaminergic neurons in the substantia nigra pars compacta (SNpc) [24]. Yet, non-motor deficits, including cognitive impairment, autonomic dysfunction and neuropsychiatric symptoms can precede the onset of motor symptoms and are highly prevalent in PD [27, 30]. Moreover, 80% of PD patients also develop dementia (PDD) after a disease duration of approximately 10-20 years [1, 50]. On the other hand, dementia with Lewy bodies (DLB), is defined by an early onset of fluctuating cognition, visual hallucinations, and dementia preceding or occurring concomitantly within one year from the onset of parkinsonism [35, 36]. PDD and DLB show considerable clinical overlap and may be considered as two ends of a disease spectrum with different timing of parkinsonism and dementia [7, 8, 44]. Despite the variation in the temporal sequence, PDD and DLB are both pathologically characterized by the presence of α-synuclein aggregates in Lewy bodies (LBs) and Lewy neurites (LNs), in the brainstem, limbic and neocortical regions [12, 36]. Of interest in this respect, the insular cortex could contribute to some of the overlapping non-motor symptoms due to its wide range of functions and connections to the rest of the brain.

The insula is involved in the integration of somatosensory and autonomic information with higher cognitive functions [5, 17, 25]. It also plays a role in emotion recognition, cognition, and awareness of interoceptive information and thus acts as the basis of self-awareness [19]. It has been identified as the neural substrate for many neuropsychiatric disorders such as anxiety, depression, and bipolar disorder [28]. According to Braak staging in PD, α-synuclein pathological aggregates progress from brainstem to limbic brain regions in the prodromal and early stage of the disease followed by the neocortex in more advanced stages. The insula is affected in advanced stages of the disease [12, 13]. Atrophy of the insula in PD, assessed by neuroimaging, correlated with executive dysfunction, one of the most common and early cognitive dysfunctions in PD [34]. Moreover, a reduction in dopaminergic receptor binding and grey matter density has been associated with mild cognitive impairment (MCI) in PD [18, 33]. Although the insula lately received much attention due to neuroimaging advances, little is known regarding its neuropathological characteristics in PD and DLB.

The insula is a heterogeneous region hidden deep within the Sylvian fissure and is widely connected to the brain. Macroscopically, the insula is divided into anterior and posterior gyri both constituting different cytoarchitectures. Microscopically and in order from ventro-rostral to dorso-caudal, the main sub-regions are defined as peri-allocortical agranular (Ia), pro-isocortical dysgranular (Id), and isocortical granular (Ig) and hypergranular (G) sub-regions based on the cytoarchitecture and number of layers [23, 26, 32, 40]. According to the location and connectivity, the agranular insula relays projections mostly to limbic regions and the granular insula projects mostly to cortical areas [5, 38]. Preferential projections to the anterior insula arise from the pre-piriform olfactory, orbitofrontal, and rhinal cortices. While only the posterior insula receives projections from the 2ry somatosensory areas along with projections from cingulate cortex [38, 41]. Thus, the transition of the insular cytoarchitecture from allocortical to isocortical represents an important feature, reflecting not only its diversity but also its preferential connections and functions. Based on the current staging criteria with earlier involvement of the allocortical limbic cortex compared to the neocortex in PD, the diversity of the insular sub-regions could provide insight into the underlying characteristics predisposing to degeneration. Moreover, the presence of von Economo neurons (VENs), spindle shaped neurons in layer Vb, mostly in the agranular insula adds to the uniqueness of the region. VENs are implied to play a role in emotion, cognition, interoception, decision-making, and social awareness [4, 16].

Considering the wide-spread connectivity of the insula, the cellular heterogeneity and differential functional properties of the insular sub-regions, we hypothesize that the allocortical sub-region of the insula displays greater vulnerability than the isocortical sub-regions in PD and DLB. To gain insight in the selective vulnerability of the insular sub-regions and its cell types, we performed a detailed analysis of the α-synuclein distribution pattern in postmortem brain tissues of subjects with incidental Lewy body disease (iLBD), PD and DLB.

## Materials & Methods

### Post-mortem human brain tissue

Insular post-mortem tissues from 8 donors with iLBD, PD(D), and DLB (age range: 67-93 years) and 2 age-matched controls (range: 70-78 years) were collected by the Netherlands Brain Bank and the department of Anatomy and Neurosciences, VU University Medical Center, Amsterdam. All donors had provided written informed consents for donation of brain tissue and access to clinical and neuropathological reports in compliance with ethical and legal guidelines. The main inclusion criteria were: 1) clinical diagnosis of PD(D) or DLB according to revised MDS diagnostic criteria [36, 45] and 2) pathological confirmation of diagnosis [2]. The iLBD donor had no record of neurological or neuropsychiatric symptoms yet neuropathological assessment revealed α-synuclein pathology in the brain. Subjects were excluded if they had a long history of neuropsychiatric disorders, or suffered from insular infarcts.

The entire insula was dissected into 0.5 cm thick slices and defined according to its borders with orbitofrontal and temporal cortices inferiorly and inferior frontal gyrus operculum superiorly. Tissue blocks were cryo-protected with 30% sucrose, frozen and stored at -30°C until further processing. The tissue slices were then sectioned using a sliding microtome into 60 μm thick sections.

### Neuropathological assessment

For neuropathological diagnosis and staging, 6μm paraffin sections from several brain regions of all donors were stained for α-synuclein, β-amyloid, hyperphosphorylated tau, hematoxylin and eosin (H&E), TDP-43, and congo-red according to current consensus diagnostic guidelines **[2]**. Confirmation of either iLBD, PD or DLB and concommitant AD pathology was based on guidelines using Braak staging for neurofibrillary tangles (Braak NFT 0-6), Braak α-synuclein (Braak α-syn 0-6), Thal phase for β-amyloid (0-5), and ABC scoring system **[10, 12, 29, 39, 51]**. Glial tauopathies such as age related tauopathy of the astroglia (ARTAG) and primary age related tauopathy (PART) were assessed primarily in the temporal cortex and amygdala in all donors **[20, 31]**. α-synuclein inclusions in the insular cortex were assessed semi-quantitatively in 60μm sections. The semi-quantitative ordinal scoring system was developed based on the recommendations of the consensus diagnostic criteria for DLB with consideration of the tissue thickness (supplementary-Table S1).

### Immunohistochemistry

For α-synuclein immunostaining, 60μm sections were pre-treated with 98% formic acid (Sigma, Steinheim, Germany) for 10 minutes at room temperature (RT). Endogenous peroxidase activity was blocked by 0.3% H2O2 in Tris-buffer saline (TBS) at RT and the sections were subsequently incubated in TBS containing 0.5% Triton-x 100 (TBS-Tx), 2% bovine serum albumin, and primary antibody mouse anti-α-synuclein (1:2000; 610786; BD Biosciences, Europe) for 24 hours at RT. Following rinsing in TBS, the sections were incubated in the 2ry antibody biotinylated horse anti-mouse IgG (1:200, Vector) in the same buffers for 2 hours followed by standard avidin-biotin complex (1:200, Vectastatin ABC kit, Standard; Vector Laboratories, California, USA) in TBS. Then 3,3′-diaminobenzidine (DAB) was used to visualize staining. The sections were mounted and counter-stained with thionin (0.13%, Sigma) for visualization of neurons and definition of sub-regions. Adjacent sections were then immunostained with an astrocytic marker. Tissues were pre-treated with citrate buffer pH 6.0 in a steamer (95^o^C) then blocked with 5% normal donkey serum in TBS (Jackson, UK) and incubated in rabbit anti-GFAP(1:10000; Z0334; DAKO, Denmark) in TBS-T for 72 hours at 4°C. Subsequently, sections were incubated in biotinylated goat anti-rabbit (1:200, DAKO) for 90 minutes followed by ABC (1:400) and DAB. For immunofluorescent double staining of α-synuclein and GFAP, sections were pre-treated with formic acid (98%) for 5 minutes then blocked with normal donkey serum (5%). Tissues were then incubated in rabbit anti-GFAP(1:4000; DAKO) and mouse anti α-synuclein (1:1000; BD Biosciences) for 72 hours at 4°C. This was followed by incubation in donkey anti-mouse coupled with Alexa Fluor 488 (1:400; Mol. Probes), donkey anti-rabbit coupled with Alexa Fluor 594, and Diamidino-2-Phenylindole(4,6)dihydrochloride (DAPI, Sigma) for 2 hours. The tissue sections were then mounted on glass slides and cover-slipped with mowiol as mounting medium (4-88 Calbiochem).

### Brightfield and confocal laser scanning microscopy

Digital images of the immuno-stained slides were made with a photomicroscope (Leica DM5000) equipped with color camera DFC450, Leica LASV4.4 software and 63x oil objective lens. Immunofluorescent labelling was visualized using confocal laser scanning microscopy LEICA TCS SP8 (Leica Microsystems). Image acquisition was done using 100x/1.4 NA objective lens, 405 nm diode, and pulsed white light laser (80Hz) with excitation wavelengths 405, 499 and 598nm. Deconvolution of image stacks was performed using Huygens Professional software (Scientific Volume Imaging, The Netherlands). Co-localization analyses to assess co-occurrence of GFAP and α-synuclein were performed using Imaris software 8.3 (Bitplane, South Windsor, CT, USA).

### Definition of insular sub-regions

The anatomical and cytoarchitectural characteristics of the insular subregions were identified in Nissl stained sections by YF and WvdB based on the granularity and density of layers II and IV. For simplicity, definitions were based on the four known sub-regions: agranular, dysgranular, and granular/hypergranular insula **[1]**. The agranular insula was defined based on its ventral anterior location, absence of layers II and IV, and clusters of VENs in layer Vb. The dysgranular region is dorsal to the agranular and has more distinguished layers II and IV. The granular and hypergranular regions were defined based on their dorso-caudal and mid to posterior location and consisted of increasingly dense and granular layers II and IV **[40]** (Fig. 1). VENs and fork cells were assessed in layer V of the agranular insula and VENs were defined based on their spindle-shaped morphology and anatomical location **[47]**.

**Fig. 1.**
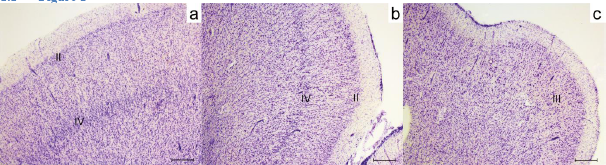
Definition of Insular sub-regions in 60 μm thick sections. (**a**) Granular insular grey matter shows uniform and well defined granular layers II and IV in an iLBD case. (**b**) Dysgranular insula shows less dense and granular layers II and IV. (**c**) Agranular insula grey matter is shown lacking layers II and IV. iLBD: incidental Lewy body disease, II: layer two, III: layer three, IV: layer four. Magnification: 25x, scale bar 500 μm

## Results

### Clinical and neuropathological characteristics of PD and DLB donors

All PD(D) and DLB donors included in this study were at moderate to severe stages of disease, clinically and neuropathologically. Braak α-synuclein stages ranged from III to VI and disease duration ranged from two to fifteen years. The PDD and DLB donors had moderate to severe cognitive impairments at time of death, REM sleep behavioral disorder (RBD), and several neuropsychiatric symptoms were reported including hallucinations, depression, and anxiety. During general neuropathological assessment, iLBD donor (iLBD-1) revealed α-synuclein pathology in brainstem and limbic regions, according to Braak α-synuclein stage IV. Moreover, ARTAG was present in PDD-1 and 2 while the latter also contained PART, both of which represent different forms of age-related tauopathy. NFT and β-amyloid plaques were also present in all cases, and were most severe in DLB. The demographics and neuropathological staging of all donors included in this study are summarized in Table 1.

**Table 1.** Subject Demographics and neuropathological staging. ABC score: AC(03), ARTAG: Aging-related tau astrogliopathy, Braak NFT:0-6, Braak a-syn: 0-6 DLB: Dementia with Lewy bodies, HC: Healthy control, iLBD: incidental Lewy body disease, LBV: Lewy body variant, NFT: neurofibrillary tangles, PART: primary age-related tauopathy, PD: parkinson’s disease, RBD: REM sleep behavioral disorder, Thal phase: 0-5, α-syn: α-synuclein,: none/absent

| # | HC-1 | HC-2 | iLBD-1 | PD-1 | PD-2 | PDD-1 | PDD-2 | PDD-3 | DLB-1 | DLB-2 |
| --- | --- | --- | --- | --- | --- | --- | --- | --- | --- | --- |
| Sex | F | M | F | M | F | M | F | M | M | M |
| Clinical Diagnosis | HC | HC | HC | PD | PD | PDD | PDD | PDD | DLB | DLB |
| Age at death (yr) | 78 | 70 | 88 | 78 | 93 | 74 | 88 | 81 | 67 | 75 |
| Pathological diagnosis | Control | Control | iLBD | PD | PD | PDD | PDD | PDD | DLB | DLB |
| Age of onset | - | - | - | 75 | 91 | 67 | 73 | 77 | 62 | 73 |
| Disease Duration | - | - | - | 3 | 2 | 7 | 15 | 4 | 5 | 2 |
| Braak α-syn | - | - | 4 | 3 | 3 | 6 | 5 | 6 | 6 | 6 |
| Braak NFT & tauopathy | - | - | 3 | 1 | 2 | 2 + ARTAG | 2 + ARTAG, PART | 3 | 5 | 3 |
| Thal phase B-amyloid | - | - | 2 | 0 | 2 | 3 | 0 | 3 | 5 | 4 |
| ABC score | - | - | A3B2C1 | A0B1C0 | A1B1C0 | A2B1C0 | A0B1C0 | A2B2C0 | A3B3C3 | A2B2C3 |
| Psychiatric symptoms | - | - | - | - | Depression | Delirium, Anxiety, RBD, Hallucinations | Depression, Panic attacks, Hallucinations | RBD, Anxiety, Depression, Disinhibition, Hallucinations | RBD, Hallucinations, Apraxia, Capgras syndrome | Paranoia, Psychosis, Hallucinations |

### Distribution pattern of α-synuclein pathology in the insular cortex of iLBD, PD and DLB

We observed a decreasing gradient of α-synuclein pathology load from the ventro-rostral agranular sub-region to the dorsal dysgranular and dorso-caudal granular sub-regions. In iLBD and PD(D), α-synuclein deposits were present in all grey matter layers of the agranular insula, whereas in the dorsal dysgranular less immunoreactivity was observed. The α-synuclein immunoreactivity was minimal or absent in the granular insula. LNs were present in all layers while LBs were predominantly found in the deep layers V and VI. In DLB cases, the gradient was less pronounced due to the presence of severe inclusions in all sub-regions and an abundance of LBs in layer V-VI with relative sparing of layer III/IV in the granular insula (Supplementary Table S1).

The iLBD insular cortex (iLBD-1), assessed at the level of mid-insula, showed few LN in both agranular and dysgranular regions and very mild immunoreactivity in the granular insula (Table S1). α-synuclein immunoreactive features consisted of dot-like deposits, few LNs, sparse LBs, and some astroglial deposits (Fig. 2 a-c). PD-1 revealed very sparse α-synuclein inclusions in all sub-regions, more pronounced in the agranular insula and mild inclusions in the white matter of granular insula (Fig. 2 d-f). PD-2 showed moderate to severe LN and few LBs in the deep layers of the agranular insula. The granular and dysgranular sub-regions contained bulgy LNs as well as a mild to moderate number of LNs and LBs, respectively (Fig. 2 g-i).

**Fig. 2.**
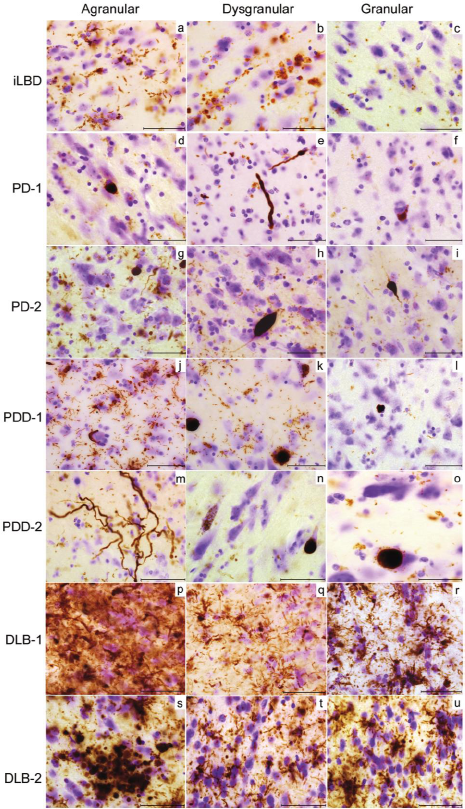
Distribution pattern of α-synuclein in insular sub-regions. iLBD shows mild LNs and astroglial α-synuclein inclusions in layer I of agranular insula (**a**), few glial inclusions in dysgranular insula (**b**), and sparse dot-like aggregates in granular insula (**c**). PD-1 agranular insula shows a LB-like inclusion and dot-like aggregates **(d),** the dysgranular insula shows bulgy LNs in layer I **(e),** and the granular insula shows an intracellular LB inclusion **(f)** PD-2 shows many LNs inclusions in agranular insula and glial α-synuclein (**g**) and less but bulgy LN in dysgranular (**h**) and granular regions (**i**). InPDD2 severe astroglial α-synuclein inclusions are shown in agranular insula (**j**) few LBs and LNs in dysgranular insula (**k**). The granular insula shows dot-like aggregates and astroglial α-synuclein **(l)**. In PDD-1 agranular insula,very long LNs and some dot-like aggregates are seen in layer I (**m**). Dysgranular insula in PDD-1 shows granular cytoplasmic inclusions in neurons and a LB **(n)** while the granular insula shows less aggregates and a LB in the infragranular layer (**o**). In DLB-1, severe α-synuclein inclusions are seen in agranular insula throughout all layers (**p**).Severe astroglial inclusions are seen in the supragranular layers of dysgranular and granular insula (**q-r**). In DLB-2, a cluster of dystrophic LNs and glial inclusions are shown in layer II of the agranular insula (**s**). The dysgranular insula contains LNs and dot-like structures (**t**) also abundant in the granular insula superficial layers (**u**). DLB: dementia with Lewy bodies, iLBD: incidental Lewy body disease, LB: Lewy bodies, LN: Lewy neurites, PD: Parkinson’s disease, PDD: parkinson’s disease dementia. Magnification: 630x, scale bar 50μm

In PDD 1-3, the same gradient of α-synuclein immunoreactivity was present with highest load of pathology in agranular insula. PDD-1 showed very long LNs in the superficial layers of the agranular insula and a mild to moderate α-synuclein load in the granular and dysgranular insula, respectively. Astroglial α-synuclein inclusions were also found in PDD 1 and 2 (Fig. 2 j-o). In contrast, DLB-1 and 2 showed severe α-synuclein inclusions in all layers and sub-regions and severe protoplasmic astroglial α-synuclein inclusions (Fig. 2 p-u).

### Morphology of α-synuclein immunoreactive structures in the insular cortex

In PDD cases with astroglial tauopathy (PDD 1 and 2), severe synucleinopathy was observed in astrocytes (Fig. 3 a-b). The supragranular layers I-III, showed moderate to severe LNs variable in shape and size, thread-like, bulgy and long. Layers V and VI contained LNs and predominance of cortical LBs, increasing in gradient from agranular to granular sub-regions. DLB insular tissues also showed astroglial α-synuclein deposits, severe load of LNs, and cortical LBs throughout multiple layers but predominantly in layers V and VI (Fig. 3-c). Astrocytes with multiple varicosities were predominantly found in the agranular and to a lesser extent in the dysgranular insula, in both controls and patients, possibly representing varicose projection astrocytes. And in PDD and DLB, degenerative changes in the form of detached astroglial processes with bulbous and doughnut-shaped end-feet were present in superficial layers. Some degenerated astrocytes in grey and white matter also contained granular accumulations along their processes, possibly representing fuzzy astrocytes (Fig. 3 d-f).

**Fig. 3.**
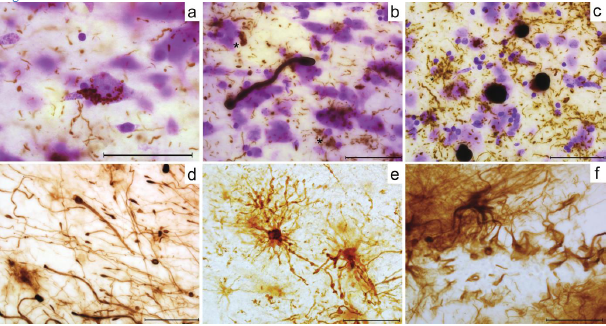
Cell specific morphology and inclusions in Lewy body diseases. In PDD-2 with astroglial tauopathy, infragranular layer of agranular insula show astroglial-to-neuronal α-synuclein inclusions (**a**), an elongated α-synuclein positive process with bulbous endings (possibly glial) surrounded by astroglial α-synuclein inclusions (**\***) and LNs (**b**). DLB-2 shows LB inclusions, LNs, and astroglial α-synuclein (**\***) within the deep infragranular layer (**c**). Loose GFAP +ve astrocytic processes are shown containing bulbuous end feet and donutshaped structures in the supragranular layers of the agranular insula in DLB-2 **(d).** PDD-1 with astroglial tauopathy shows small and dysmorphic astrocytes containing multiple varicosities within their processes, possibly fuzzy astrocytes **(e).** A GFAP +ve astrocyte is shown surrounded by unorganized processes in DLB-2 **(f).** Magnification: 630x, scale bar 50μm

### Neuronal vulnerability in insular cortex

Only 1-2% of VENs and few fork cells in layer V of fronto-insular region revealed α-synuclein inclusions in PD-2 and PDD 1 and 2. VENs showed granular cytoplasmic α-synuclein inclusions and Lewy bodies. They were also frequently surrounded by astroglial cells showing thorn-shaped α-synuclein immunoreactivity. In DLB, VENs were surrounded by inclusions but the precise location of these inclusions, intracellular or extracellular, could not be discerned due to the severe extent of pathology within the agranular sub-region (Fig. 4).

**Fig. 4.**
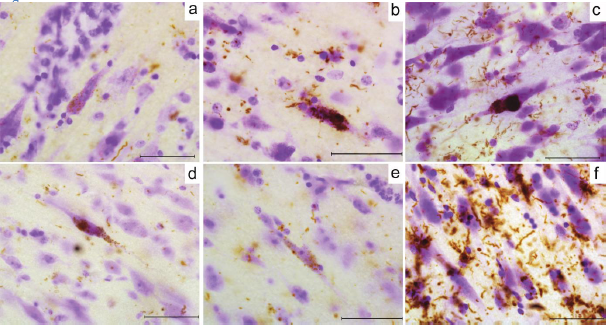
α-synuclein deposits in VEN and fork cell. (**a**) PD-2 shows granular LB inclusions (brown) along a VEN. (**b**) LB in VEN and surrounding astrocytes in PD-2. **(c)** PDD2 shows a VEN containing a large LB and multiple granular inclusions within the cell body, astroglial α-synuclein inclusions are also seen. (**d**) PDD-1 shows LB in the soma and dendrite of a VEN. (**e**) α-synuclein inclusions are shown in a fork cell in PDD-1. (**f**) DLB-2 agranular region shows many deposits surrounding and inflicting pyramidal neurons and rod shaped VEN. Magnification: 630x, scale bar 50μm

### Relationship between astrocytes and α-synuclein pathology

Double labelling of GFAP and α-synuclein was performed to examine the relationship between astrocytes and α-synuclein inclusions. A varicose projection astrocyte (VPA) in the infragranular layer in PD-2 was found containing a cluster of cytoplasmic α-synuclein forming a mesh-like structure (Fig. 5-a). Further colocalization analysis in all cases assessing the co-occurrence and correlation of α-synuclein and GFAP in VPA showed an inverse correlation, using Pearson’s correlation coefficient (PCC), in colocalized volume (PCC: -0.09). While Mander’s overlap coefficient (MOC) showed a large correlation between α-synuclein and GFAP (MOC= 0.93 and 0.27). The 3D reconstruction indicated possible compartmentalization of the α-synuclein within the astrocytic cell body (Fig. S1). Other protoplasmic and interlaminar astrocytes examined in PDD and DLB showed α-synuclein deposits around the cell body and processes (Fig. 5-e) and a large correlation between GFAP and α-synuclein was observed using MOC (Fig. S1).

**Fig. 5.**
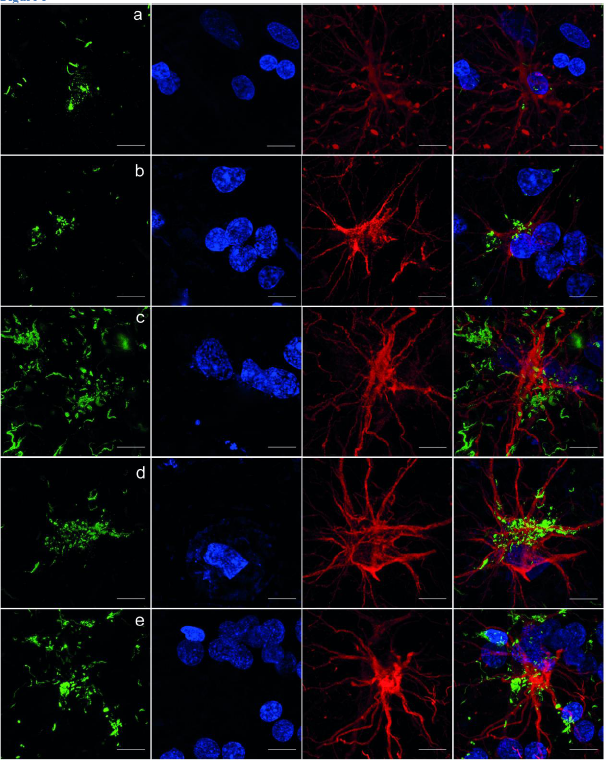
Relationship between α-synuclein immunoreactivity and astrocytes in insular cortex in PD(D) and DLB. α-synuclein (green) is present within varicose projection astrocyte (GFAP, red) cell body in infragranular layer, anterior insula in PD-2 (**a**). In PDD-2, an astrocyte is shown containing α-synuclein aggregates and surrounded by a cluster of nuclei, probably neuronal, in the anterior insula (**b**). And in PDD-3, a protoplasmic astrocyte is shown surrounded by multiple α-synuclein aggregates (**c**). DLB-1 shows α-synuclein deposits surrounding the cell body of an interlaminar astrocyte in layer I (**d**) and similar inclusions are shown within a protoplasmic astrocyte, its processes, and the surrounding clustered nuclei in posterior insula (**e**). Magnification 100x. Scale bar: 10μm

## Discussion

In this case series, we observed a decreasing gradient in the load of α-synuclein immunoreactivity from the anterior periallocortical agranular sub-region to the intermediate pro-isocortical dysgranular, and posterior isocortical granular insula of iLBD, PD and DLB subjects. This was particularly evident in iLBD and PD(D) with highest load of neuropathological inclusions in the agranular insula. In DLB, there was also extensive α-synuclein immunoreactivity in the granular region with an abundance of LBs in the infragranular layers V/VI, glial inclusions, and relative sparing of layers III/IV. Some VENs in the anterior insula revealed α-synuclein inclusions in PD(D). Astrocytes were also vulnerable to α-synuclein inclusions and showed degenerative changes at all disease stages yet most prominent in PDD and DLB. The agranular and to a lesser extent, dysgranular insula in all donors revealed varicose projection astrocytes.

The presence of a gradient across the insula from the agranular to more granular sub-regions has been previously documented for NFT and β-amyloid pathology in AD [9]. This decreasing pathological gradient appears to be consistent with differences in terms of cytoarchitecture, cell types, and myelination in the insula. Accordingly, the agranular insula comprises the highest density of acetylcholinesterase and lowest density of myelinated fibers while the opposite exists in the granular sub-regions [37]. The vulnerability of the agranular insula relative to the late and sparse involvement of the granular corresponds with an inverse relationship between myelination and neuropathological lesions in both AD and PD [14]. The agranular insula also comprises of preferential connections to the olfactory and rhinal cortices which are affected in early stage PD and show a similar allocortical cytoarchitecture [13, 38]. In line with this, the granular insula connects to isocortical somatosensory and cingulate cortex and is affected in later stages of the disease [38]. Thus the cytoarchitecture and the connectivity of the insular subregions could provide a framework for the understanding of dissemination of α-synuclein pathology throughout the brain [11, 21]. Considering the anterior insula connectivity, α-synuclein pathology and cell death in the agranular insula may also contribute to autonomic, cognitive and psychiatric symptoms in PD(D) and DLB [52].

According to the Braak staging criteria for PD, the progression of α-synuclein pathology follows a predictable pattern. Initially, α-synuclein inclusions start in the brainstem and spread caudally in the mesencephalon in stages I-III, to the mesocortex in stage IV while increasing in density and spreading to the neocortex in stages V and VI [12]. Considering the insular phylogenetic and ontogenetic variations [6], the insular sub-regions alone could reflect the global regional involvement in PD as described by Braak and colleagues. For instance, the agranular insula was shown to contain neuropathological inclusions from stage III which increased with advanced stages. While the granular insula showed mostly absent to mild inclusions at stages III&IV, and a mild to moderate load of LNs and LBs in the deep layers in stages V&VI in PDD. Yet, the pathological assessment of the insula in PD is not routinely included in all scoring criteria and is not assessed on a sub-regional basis [3]. Taking into account its allocortical to isocortical transition, assessment of the insular sub-regions could provide us with insight into disease progression in PD and DLB.

Furthermore, in our study, the granular insula in DLB showed a greater load of α-synuclein deposits compared to PDD and regardless of disease stage. Despite the greater vulnerability of the granular insula to α-synuclein deposits, relative sparing of layers III/IV was observed. The reduced vulnerability of these layers could be related to their further development in granular insula compared to other subregions, with an increasing number of neurons, parvalbumin-positive interneurons, and myelinated fibers [40]. Along with a shorter disease duration and early onset of cognitive/ neuropsychiatric deficits, DLB brains showed higher NFT and β-amyloid staging along with an extensively greater load of α-synuclein pathology in all regions including granular insula. We therefore highlight the higher vulnerability of neocortical regions, as represented by the granular insula, in DLB compared to PDD. This is important as both DLB and PDD overlap clinically and neuropathologically with an earlier onset of dementia in DLB.

We also show that VENs are vulnerable to α-synuclein pathology in advanced PD(D). VENs have been implicated in consciousness, interoception, emotion, cognition, and social awareness [4, 22, 47]. In this study, about 1-2% of VENs showed α-synuclein inclusions compared to the greater involvement of pyramidal neurons within the agranular insula. Previous studies assessing VENs found tau inclusions as well as significant cell loss in Pick’s disease compared to AD [46]. VENs are projection neurons with a uniquely and sparsely branched basal dendrites. They are known to be selectively vulnerable to degeneration early in frontotemporal dementia as well as early onset schizophrenia [26]. Despite previous implications on the possible role of VENs in PD, our study is the first to show their involvement in PDD and DLB. The VENs unique properties, location, and functions could help elucidate their role in the development of cognitive and neuropsychiatric deficits in PDD and DLB.

Astrocytes have been previously shown to contain α-synuclein inclusions in advanced PD and parallel to the neuronal involvement [15]. However, minimal astrocytic activation and cytoplasmic α-synuclein inclusions were observed in PD compared to other neurodegenerative diseases [49]. In our series, we observed the enwrapment of astrocytic cell bodies and processes with α-synuclein. Protoplasmic and interlaminar astrocytes showed extensive synucleinopathy most severe in the agranular insula, particularly in PDD with glial tauopathy and DLB. Astroglial tauopathy could play an important role by functioning as a potential seed for glial synucleinopathy and leading to further degeneration by promoting their oligomerization and coaggregation [48]. Yet another unique feature in the agranular insula, was the presence of varicose projection astrocytes (VPA). These recently discovered astrocytes with varicosities along their processes were found only in higher order primates and humans [43]. Recent studies proposed that they may provide alternative pathways for long distance communication through cortical layers [42, 43]. We report a selective topographic distribution of these cells with predominance in agranular insula and development of intracellular inclusions in PD.

The limitations of this study include a small sample size, limited range of neuropathological stages and lack of longitudinal clinical data on the cognitive and psychiatric performance of the included PD and DLB patients. Future studies including a large number of cases and quantitative assessment of related proteinopathies could warrant a better understanding of the selective vulnerability of different cell types and their connections in the insular sub-regions in neurodegenerative diseases. Moreover, assessment of the relationship between insular pathology, cognitive phenotypes and psychiatric disturbances in PD and DLB could also provide further clues towards the substrates of cognitive and neuropsychiatric deficits in these disorders.

## Conclusions

In conclusion, the distribution pattern of α-synuclein pathology revealed a decreasing anterior-to-posterior gradient in the insular cortex representative of the differential cytoarchitectural vulnerability in PD. α-synuclein pathology was more pronounced in the posterior isocortical insula in DLB compared to PDD and regardless of disease stage, thus portraying differences between these two entities. Yet, DLB and PDD cases that presented with astroglial tauopathy shared a similar astrocytic involvement and extensive astroglial synucleinopathy. We also show, for the first time, the vulnerability of VENs to α-synuclein pathology which may in turn contribute to cognitive and neuropsychiatric deficits in theses diseases. Considering the widespread insular connectivity and functions, we propose future considerations of the insular subregions for neuropathological assessment and for differential postmortem diagnosis of PD(D) and DLB.

## Abbreviations

AD: Alzheimer’s disease
ARTAG: Age related tauopathy of the astroglia
DLB: Dementia with Lewy bodies
GFAP: gial fibrillary acidic protein
iLBD: incidental Lewy body disease
LB: Lewy bodies
LN: Lewy neurites
MOC: Mander’s overlap coefficient
NFT: Neurofibrillary tangles
PART: primary age related tauopathy
PCC: Pearson’s correlation coefficient
PD: Parkinson’s disease
PDD: Parkinson’s disease with dementia
RBD: REM sleep behavioral disorder
VENs: von Economo neurons
VPA: varicose projection astrocytes

## Acknowledgments

This work was funded by Stichting ParkinsonFonds, the Netherlands.

The authors are thankful to all donors and their families for making this research possible. We would like to express our gratitude for the efforts of the Netherlands Brain Bank team for providing the necessary material and data. Finally, we would like to thank Evelien T. Huisman, Lucienne te-Bulte Baks, Allert J. Jonker, and John Bol for technical assistance.

## Declarations

The authors declare having no competing interests.

## Author Contributions

The study design was carried out by WB and YF. YF carried out the experimental work and wrote the initial manuscript. WB and AR performed the autopsies and the diagnostic scoring. AD provided technical advice and revised the manuscript. FJ participated in the design and revised the manuscript. Significant contributions were provided by WB, AD, AR, and FJ and the manuscript was then edited and finalized by YF and WB.

